# Whole-cell segmentation of tissue images with human-level performance using large-scale data annotation and deep learning

**DOI:** 10.1101/2021.03.01.431313

**Authors:** Noah F. Greenwald, Geneva Miller, Erick Moen, Alex Kong, Adam Kagel, Christine Camacho Fullaway, Brianna J. McIntosh, Ke Leow, Morgan Sarah Schwartz, Thomas Dougherty, Cole Pavelchek, Sunny Cui, Isabella Camplisson, Omer Bar-Tal, Jaiveer Singh, Mara Fong, Gautam Chaudhry, Zion Abraham, Jackson Moseley, Shiri Warshawsky, Erin Soon, Shirley Greenbaum, Tyler Risom, Travis Hollmann, Leeat Keren, Will Graf, Michael Angelo, David Van Valen

## Abstract

Understanding the spatial organization of tissues is of critical importance for both basic and translational research. While recent advances in tissue imaging are opening an exciting new window into the biology of human tissues, interpreting the data that they create is a significant computational challenge. Cell segmentation, the task of uniquely identifying each cell in an image, remains a substantial barrier for tissue imaging, as existing approaches are inaccurate or require a substantial amount of manual curation to yield useful results. Here, we addressed the problem of cell segmentation in tissue imaging data through large-scale data annotation and deep learning. We constructed TissueNet, an image dataset containing >1 million paired whole-cell and nuclear annotations for tissue images from nine organs and six imaging platforms. We created Mesmer, a deep learning-enabled segmentation algorithm trained on TissueNet that performs nuclear and whole-cell segmentation in tissue imaging data. We demonstrated that Mesmer has better speed and accuracy than previous methods, generalizes to the full diversity of tissue types and imaging platforms in TissueNet, and achieves human-level performance for whole-cell segmentation. Mesmer enabled the automated extraction of key cellular features, such as subcellular localization of protein signal, which was challenging with previous approaches. We further showed that Mesmer could be adapted to harness cell lineage information present in highly multiplexed datasets. We used this enhanced version to quantify cell morphology changes during human gestation. All underlying code and models are released with permissive licenses as a community resource.

## Introduction

Understanding the structural and functional relationships present within tissues is a challenge that is at the forefront of basic and translational research. Recent advances in multiplexed imaging have dramatically expanded the number of transcripts and proteins that can be quantified in a single tissue section while also improving the throughput of these platforms^1–12^. These technological improvements have opened up exciting new frontiers for large-scale analysis of human tissue samples. Ambitious collaborative efforts such as the Human Tumor Atlas Network^13^, the Human BioMolecular Atlas Program^14^, and the Human Cell Atlas^15^ are now using novel imaging techniques to comprehensively characterize the location, function, and phenotype of the cells in the human body. By generating high-quality, open-source datasets characterizing the full breadth of human tissues, these datasets will be as transformative as the Human Genome Project in unleashing the next era of biological discovery.

Despite this immense promise, the tools to facilitate the analysis and interpretation of these datasets at scale do not yet exist. The clearest example of this shortcoming is the lack of a generalized algorithm for locating single cells in images. Unlike flow cytometry or single-cell RNA sequencing methods, in which individual cells are dissociated and physically separated from one another prior to being analyzed, tissue imaging is performed with intact specimens. Thus, in order to extract single-cell information from images, each pixel must be assigned to a cell after image acquisition in a process known as cell segmentation. Since the features extracted through this process are the basis for downstream analyses like cell-type identification and tissue neighborhood analyses^16^, inaccuracies at this stage have far-reaching consequences for interpreting image data.

Achieving accurate and automated cell segmentation for tissues remains a substantial challenge. Depending on the tissue, cells can be rare and dispersed within a large bed of extracellular matrix or densely packed such that contrast between adjacent neighbors is limited. Cell size in non-neuronal mammalian tissues can vary over two orders of magnitude^17^, while cell morphology can vary widely from small mature lymphocytes with little discernible cytoplasm, to elongated spindle-shaped fibroblasts, to large multinucleated osteoclasts and megakaryocytes^18^. Achieving accurate cell segmentation has been a long-standing goal of the biological image analysis community, and a diverse array of software tools have been developed to meet this challenge^19–24^. While these efforts have been crucial for advancing our understanding of biology across a wide range of domains, they fall short for tissue imaging data. A common shortcoming has been the need to perform manual, image-specific adjustments to produce useful segmentations. This lack of full automation poses a prohibitive barrier given the increasing scale of tissue imaging experiments.

Recent advances in deep learning have transformed the field of computer vision, and are increasingly being used for a variety of tasks in biological image analysis, including cell segmentation^25–31^. These methods differ from conventional algorithms in that they learn how to perform tasks from annotated data. While the accuracy of these new, data-driven algorithms can render difficult analyses routine, using them in practice can be challenging: high accuracy requires a substantial amount of annotated data. Generating ground-truth data for cell segmentation is time intensive due to the need to generate pixel-level labels; as a result, existing datasets are of modest size (10^4^-10^5^ annotations). Moreover, most public datasets^26,27,32–38^ annotate the location of cell nuclei rather than the whole cell. Deploying pre-trained models to the life science community is also difficult, and has been the focus of a number of recent works^39–42^. Despite deep learning’s potential, these challenges have caused whole-cell segmentation in tissue imaging data to remain an open problem.

Here, we sought to close these gaps by creating an automated, simple, and scalable algorithm for nuclear and whole-cell segmentation that performs accurately across a diverse range of tissue types and imaging platforms. Developing this algorithm required two innovations: (1) a scalable approach for generating large volumes of pixel-level training data in tissue images and (2) an integrated deep learning pipeline that utilizes these data to achieve human-level performance. To address the first challenge, we developed a crowdsourced, human-in-the-loop approach for segmenting cells in tissue images where humans and algorithms work in tandem to produce accurate annotations (**Figure 1a**). We used this pipeline to create TissueNet, a comprehensive segmentation dataset of >1 million paired whole-cell and nuclear annotations. These curated annotations were derived from images of nine different organs acquired from six distinct imaging platforms. TissueNet is the largest cell-segmentation dataset assembled to date, containing twice as many nuclear and 16 times as many whole-cell labels as all previously published datasets combined. To address the second challenge, we developed Mesmer, a deep learning pipeline for scalable, user-friendly segmentation of imaging data. Mesmer was trained on TissueNet and is the first algorithm to demonstrate human-level performance on cell segmentation. To enable broad use by the scientific community, we harnessed DeepCell, an open-source collection of software libraries, to create cloud-native software for using Mesmer, including plugins for ImageJ and QuPath. We have made all underlying code and trained models available under a permissive license as a community resource, setting the stage for application of these modern, data-driven methods to a broad range of fundamental and translational research challenges.

**Figure 1:**
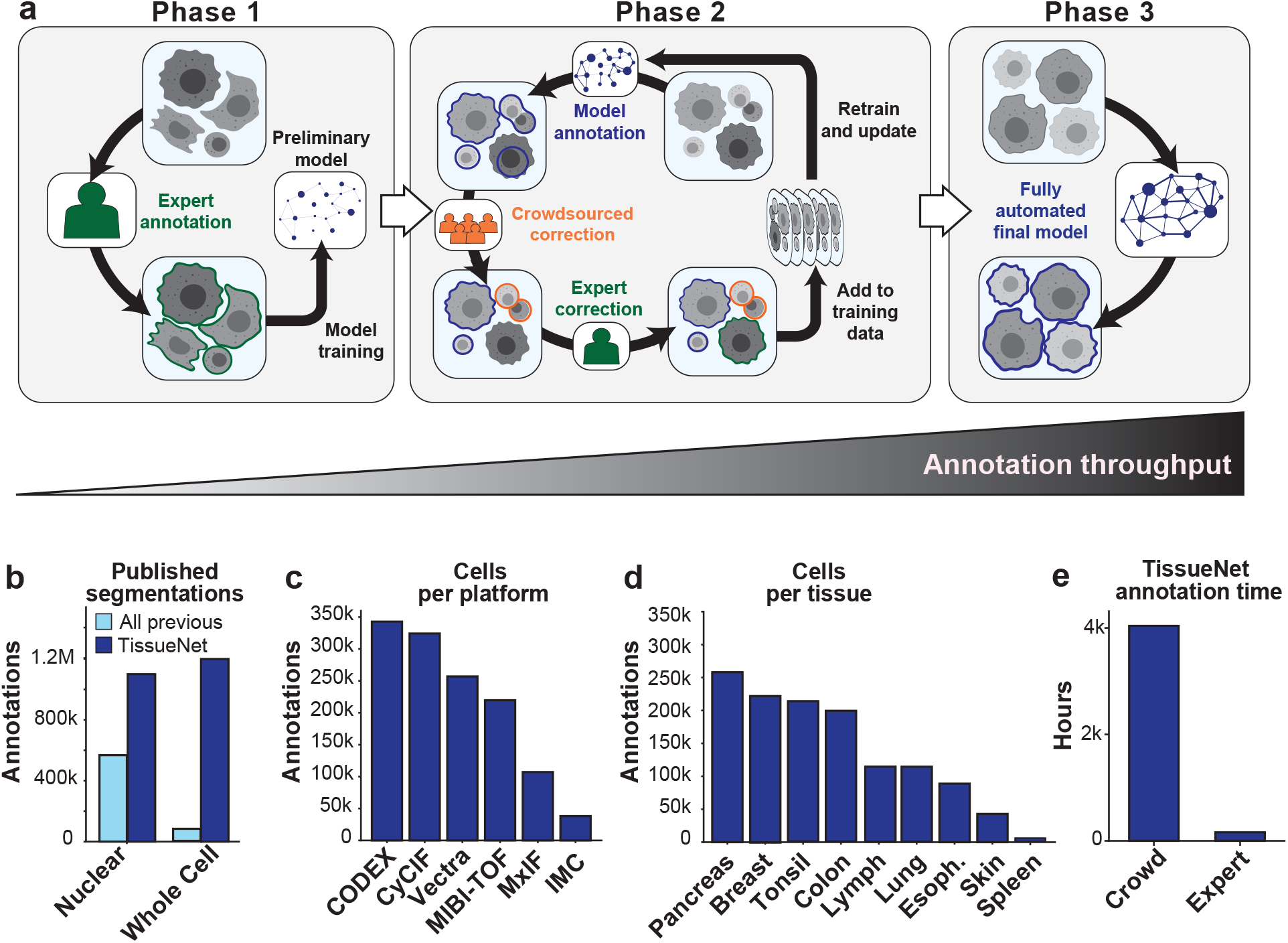
A human-in-the-loop approach enables scalable, pixel-level annotation of large image collections. **a**, This approach has three phases. During phase 1, annotations are created from scratch to train a model. During phase 2, new data are fed through a preliminary model to generate predictions. These predictions are used as a starting point for annotators to correct. As more images are corrected, the model improves, which decreases the number of errors, increasing the speed with which new data can be annotated. During phase 3, an accurate model is run without human correction. **b**, TissueNet has more nuclear and whole-cell annotations than all previously published datasets. **c**, The number of cell annotations per platform in TissueNet. **d**, The number of cell annotations per tissue type in TissueNet. **e**, The number of hours of annotation time required to create TissueNet.

### A human-in-the-loop approach drives scalable construction of TissueNet

Existing annotated datasets for cell segmentation are limited in scope and scale (**Figure 1b**)^26,27,32–38^. This limitation is largely due to the linear, time-intensive approach used to construct them, which requires the border of every cell in an image to be manually demarcated. This approach scales poorly, as the time required to label each image remains constant throughout the annotation effort. We therefore implemented a three-phase approach to create TissueNet. In the first phase, expert human annotators outlined the border of each cell in 80 images. The labeled images were then used to train a preliminary model (**Figure 1a, left;** Methods). Once the preliminary model reached a sufficient level of accuracy, correcting mistakes required less time than labeling from scratch. Although the exact point at which this transition occurs depends on model quality and training data diversity, we found that 10,000 cells was a reasonable approximation.

The process then moved to the second phase **(Figure 1a, middle)**, where images were first passed through the model to generate predicted annotations. These predictions were sent to crowdsourced annotators to correct errors. The corrected annotations then underwent final inspection by an expert prior to being added to the training dataset. When enough new data were compiled, a new model was trained and phase two was repeated. Each iteration yielded more training data, which led to improved model accuracy and fewer errors that needed to be manually corrected. This virtuous cycle continued until the model achieved human-level performance. At this point, we transitioned to the third phase (**Figure 1a, right**), where the model was run without human assistance to produce high-quality predictions. One advantage of this approach is that we utilized annotators with different amounts of bandwidth and expertise: experts have experience but limited bandwidth, while crowdsourced annotators have limited experience but higher bandwidth. Triaging each task according to its difficulty and accessing a much larger pool of human annotators further reduced the time and cost of dataset construction.

Human-in-the-loop pipelines require specialized software that is optimized for the task and can be scalably deployed. We therefore developed DeepCell Label^43^, a browser-based graphical user interface optimized for editing existing cell annotations in tissue images (**Figure S1a**, Methods). DeepCell Label is supported by a scalable cloud backend that dynamically adjusts the number of servers according to demand (**Figure S1b**). Using DeepCell Label, we trained annotators from multiple crowdsourcing platforms to identify whole-cell and nuclear boundaries. To further simplify our annotation workflow, we integrated DeepCell Label into a pipeline that allowed us to prepare and submit images for annotation, have users annotate those images, and download the results. The images and resulting labels were used to train and update our model, completing the loop (**Figure S1c;** Methods).

Our goal in creating TissueNet was to use it to power general-purpose tissue segmentation models. To ensure that models trained on TissueNet would serve as much of the imaging community as possible, we made two key choices. First, all data in TissueNet contains two channels, a nuclear channel (such as DAPI) and a membrane or cytoplasm channel (such as E-cadherin or Pan-Keratin). Although some highly multiplexed platforms are capable of imaging dozens of markers at once^1,2,4,6^, restricting TissueNet to include only the minimum number of channels necessary for whole-cell segmentation maximizes the number of imaging platforms where the resulting models can be used. Second, the data in TissueNet are derived from a wide variety of tissue types, disease states, and imaging platforms. This diversity of data allows models trained on TissueNet to handle data from many different experimental setups and biological questions. The images included in TissueNet were acquired from the published and unpublished works of labs who routinely perform tissue imaging^44–51^. Thus, while this first release of TissueNet encompasses the tissue types most commonly analyzed by the community, we expect that subsequent versions of TissueNet will be expanded to include less-studied organs.

As a result of the scalability of our human-in-the-loop approach to data labeling, TissueNet is larger than the sum total of all previously published datasets^26,27,32–38^ (**Figure 1b**), with 1.3 million whole-cell annotations and 1.2 million nuclear annotations. TissueNet contains data from six imaging platforms (**Figure 1c)**, nine organs (**Figure 1d**), and includes both histologically normal and diseased tissue (e.g., tumor resections). TissueNet also encompasses three species, with images from human, mouse, and macaque. Constructing TissueNet required >4,000 person hours, the equivalent of nearly 2 person-years of full-time effort (**Figure 1e**). With an average hourly rate of $6 per hour, we anticipate that subsequent datasets of this size will cost around USD $25,000 to produce—a significant reduction versus highly trained ($30/h) or expert pathologist (>$150/h) annotators.

### Mesmer is a novel algorithm for accurate whole-cell segmentation of tissue data

An ideal deep learning model for cell segmentation has two specific requirements. First, a suitable model must be accurate, which is challenging given the range of cell morphologies, tissue types, and image platforms present in TissueNet. A model capable of accurately performing whole-cell segmentation in this setting needs sufficient representational capacity to understand and interpret these heterogeneous images. Second, a suitable model needs to be fast. Image datasets are increasing rapidly in size, and a model with high performance but poor inference speed would be of limited utility.

To satisfy these requirements we developed the PanopticNet deep learning architecture. To ensure adequate model capacity, PanopticNets use a ResNet50 backbone coupled to a modified Feature Pyramid Network (FPN)^52–54^ (**Figure S2a**; Methods). ResNet backbones are a popular architecture for extracting features from imaging data for a variety of tasks^54^. FPNs aggregate features across length scales, producing representations that contain both low-level details and high-level semantics^52^. To perform segmentation, two semantic heads are attached to the highest level of the FPN to create pixel-level predictions. These heads perform two separate prediction tasks. The first head predicts whether a pixel is inside a cell, at the cell boundary, or part of the image background^25,26^. The second head predicts the distance of each pixel within a cell to the cell centroid (**Figure S2a**; Methods); we extended previous work^30,55^ by explicitly accounting for cell size in this step.

We used the PanopticNet architecture and TissueNet to create Mesmer, a deep learning pipeline for accurate nuclear and whole-cell segmentation of tissue data. Mesmer’s PanopticNet model contains four semantic heads (two for nuclear segmentation and two for whole-cell segmentation) that are attached to a common backbone and FPN. The input to Mesmer is a nuclear image (e.g. DAPI) to define the nucleus of each cell and a membrane or cytoplasm image (e.g. CD45 or E-cadherin) to define the shape of each cell (**Figure 2a**). These inputs are normalized^56^ (to improve robustness), tiled into patches of fixed size (to allow processing of images with arbitrary dimensions), and then fed to the PanopticNet model. The model outputs are then untiled^57^ to produce predictions for the centroid and boundary of every nucleus and cell in the image. The centroid and boundary predictions are used as inputs to a watershed algorithm^58^ to create the final instance segmentation mask for each nucleus and each cell in the image (Methods).

**Figure 2:**
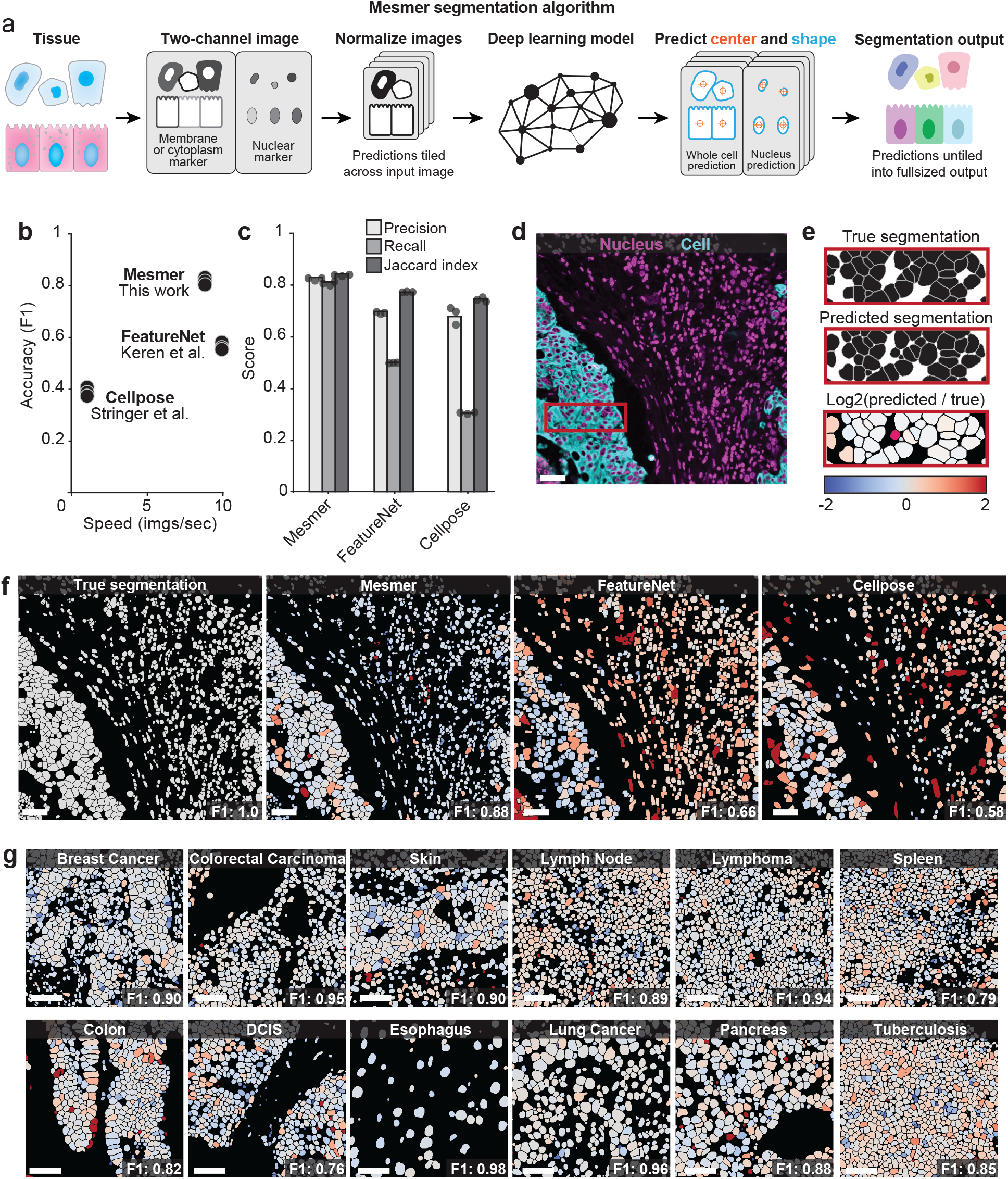
Mesmer delivers accurate nuclear and whole-cell segmentation in multiplexed images of tissues. **a**, Diagram illustrating the key steps in the Mesmer segmentation pipeline. **b**, Speed versus accuracy comparison of Mesmer and previously published models, assessed on the test set of TissueNet. Accuracy is measured by the F1 score (Methods) between the predicted segmentations and the ground-truth labels, where 0 indicates no agreement and 1 indicates perfect agreement. **c**, Evaluation of precision, recall, and Jaccard index demonstrates that Mesmer outperforms previously published models. **d**, Color overlay of representative image of colorectal carcinoma. **e**, Inset showing the ground truth (top) and predicted (middle) labels from a small region in **d**, along with a visual representation of segmentation accuracy (bottom). Predicted segmentations for each cell are colored by the log_2_ of the ratio between the predicted area and ground-truth area. Predicted cells that are too large are red, while predicted cells that are too small are blue. **f**, Ground-truth segmentation labels for the image in **d**, along with the predicted labels from Mesmer and previously published models, each colored by the log_2_ as in **e**. As seen visually, Mesmer offers substantially improved performance in comparison to previous methods. **g**, Mesmer generalizes across tissue types, imaging platforms, and disease states. The F1 score is given for each image. In all panels, scale bars are 50 μM.

We used the newly created TissueNet dataset to train Mesmer’s model. We randomly partitioned TissueNet into training (80%), validation (10%), and testing (10%) splits. The training split was used to directly update the model weights during training, with the validation split used to assess increases in model accuracy after each epoch. The test split was completely held out during training and used only to evaluate model performance after training. We used standard image augmentation during training to increase model robustness. To benchmark model accuracy, we built off our prior framework for classifying segmentation errors^37^. In brief, we perform a linear assignment between predicted cells and ground truth cells. Cells that map 1-to-1 with a ground truth cell are marked as accurately segmented; all other cells are assigned to one of several error modes depending on their relationship with the ground truth data. We use these assignments to calculate precision, recall, F1 score, and Jaccard index; see the Methods section for detailed descriptions.

We benchmarked Mesmer against two previously published algorithms for segmenting tissue image data. The first was our own model for tissue segmentation, FeatureNet^26^, which we previously used^16^ to perform nuclear segmentation followed by expansion to analyze a cohort of MIBI-TOF breast cancer samples. The second was Cellpose^28^, a recently published algorithm for whole-cell segmentation of microscopy data. Overall, we observed higher accuracy for Mesmer (F1=0.82) than both FeatureNet (F1=0.63) and Cellpose (F1=0.41) (**Figure 2b**). Mesmer is only 13% slower than FeatureNet, despite a significant increase in model capacity, and is 20 times faster than Cellpose (**Figure 2b**). This speed advantage is due primarily to differences in post-processing between the two approaches, which accounts for the majority of computational time (**Figure S2b**). In addition, Mesmer has higher recall (% of true cells correctly identified), precision (% of predictions that are correct), and Jaccard index (% overlap between predicted and true segmentations) than both FeatureNet and Cellpose (**Figure 2c**). Finally, we compared Mesmer to alternative architectures^59^ that we trained on TissueNet in order to understand the relative contributions of the PanopticNet architecture and the TissueNet dataset to performance. We also compared Mesmer with published, pre-trained nuclear segmentation models^60^; we performed a pixel expansion (the standard approach^16,50,61,62^ to approximate the entire cell) on the outputs of these models to enable whole-cell benchmarking (**Figure S2c-g**). In all cases, Mesmer outperformed existing methods, demonstrating the advantages of Mesmer’s whole cell segmentation approach.

To visualize the performance differences of the three models, we segmented an image of colorectal carcinoma with Mesmer, FeatureNet, and Cellpose (**Figure 2d**). We compared segmentation predictions to the ground-truth data, coloring each cell by the ratio of the predicted area to the ground-truth area (**Figure 2e**). Overall, Mesmer more effectively captured the true size of each cell in the image (**Figure 2f**). In comparison, FeatureNet poorly captured the true size of each cell, which is expected given that this model approximates cell shape by performing nuclear segmentation followed by expansion. In line with its lower recall score (**Figure 2c**), Cellpose failed to identify a large fraction of the cells in the image (**Figure 2f**), likely due to the relative scarcity of tissue images in the data used to train Cellpose.

Finally, we examined Mesmer’s segmentation predictions across a range of tissue types (**Figure 2g**). Importantly, Mesmer’s errors were unbiased, with an equal number of cells that were too large or too small. Further, Mesmer’s errors were not correlated with the true size of the cell (**Figure S2h**). In contrast, methods that rely on nuclear segmentation and expansion tend to overestimate the size of most small cells and underestimate the size of most large cells (**Figure S2h**). Taken together, this benchmarking demonstrates that Mesmer is a significant advance over prior segmentation methods.

### Mesmer achieves human-level performance for whole-cell segmentation

The overall goal of this work was to create a general-purpose model that would perform well across all tissue types and imaging platforms in TissueNet without case-by-case optimization. Our results thus far (**Figure 2g, Figure 3a**) suggested that Mesmer achieved this goal. However, tissue structure and cell morphology can vary widely depending on organ site and diagnosis^17,18^. This variation is further complicated by platform-specific differences in signal-to-noise ratio, imaging resolution, autofluorescence, and background staining. Therefore, training a specialist model on data from a single platform or single tissue type may lead to superior performance on that data when compared to a model trained on all of the data in TissueNet (e.g. Mesmer’s model).

**Figure 3:**
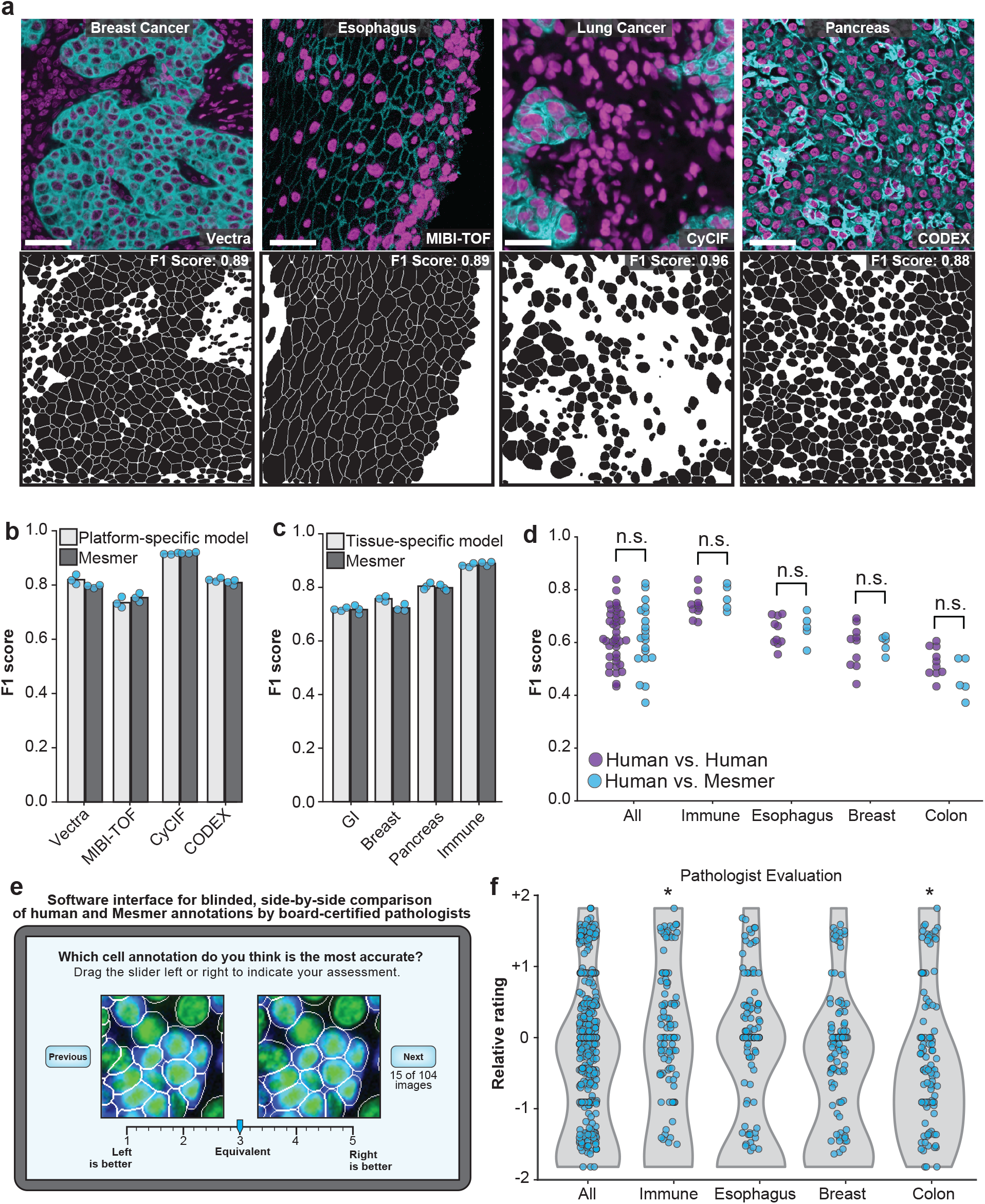
Mesmer performs whole-cell segmentation across tissue types and imaging platforms with human-level accuracy. **a**, Sample images, predicted segmentations, and F1 scores for distinct tissues and imaging platforms visually demonstrate that Mesmer delivers accurate cell segmentation for all available imaging platforms. **b**, Mesmer has equivalent accuracy to specialist models trained only on data from a specific imaging platform (Methods), with all models evaluated on data from the platform used for training. **c**, Mesmer has equivalent accuracy to specialist models trained only on data from a specific tissue type (Methods), with all models evaluated on data from the tissue type used for training. **d**, F1 scores evaluating the agreement between segmentation predictions for the same set of images. The predictions from five independent expert annotators were compared against each other (human vs. human) or against Mesmer (human vs. Mesmer). No statistically significant differences between these two comparisons were found, demonstrating that Mesmer achieves human-level performance. **e**, Workflow for pathologists to rate the segmentation accuracy of Mesmer compared with expert annotators. **f**, Pathologist scores from the blinded comparison. A positive score indicates a preference for Mesmer while a negative score indicates a preference for human annotations. Pathologists displayed no significant preference for human labels or Mesmer’s outputs, again demonstrating that Mesmer has achieved human-level performance. n.s., not significant; *p<0.05, two-sample t-test. All scale bars are 50 μM.

To evaluate Mesmer’s generalizability, we benchmarked performance against models that were trained using a subset of TissueNet that was either tissue- or platform-specific. We observed comparable performance between Mesmer and the specialist models. In some cases, specialist models exhibited slightly superior performance to Mesmer (**Figure 3b, c**). However, the magnitude of these differences was modest (+/- 0.03 mean F1 score), indicating a reasonable tradeoff between accuracy and generalizability for TissueNet. We next sought to evaluate how specialist models performed when evaluated on data not seen during training. Models trained on data from immunofluorescence-based imaging platforms were relatively interchangeable (**Figure S3a**), reflecting the similarity between these imaging methods. However, models trained on immunofluorescence images did not perform well when evaluated on data from mass spectrometry-based platforms, and vice versa (**Figure S3a**). In contrast to platform-specific models, tissue-specific models had poor performance when evaluated on tissue types not seen during training (**Figure S3b**). This analysis highlights the importance of dataset diversity for producing performant deep learning models for cell segmentation.

We next sought to determine how the size of TissueNet contributed to Mesmer’s performance. We trained replicate models on increasingly large fractions of TissueNet and evaluated the marginal benefit of each additional training example. As expected, additional data made the largest difference when datasets were small; as the size of the dataset increased, the benefit from additional examples began to plateau (**Figure S3c**). This plateau occurred at a low dataset fraction for the overall F1 score, which is dominated by the easiest examples. However, several error types (such as catastrophes and merges; Methods) plateaued at a higher dataset fraction, reflecting the value of dataset scale to model performance (**Figure S3d-h**). All of these findings were contingent on having sufficiently diverse training data, as even large datasets containing only a single tissue type resulted in models that generalized poorly (**Figure S3b**).

The metrics for model accuracy used here treated human annotated data as ground truth. However, given the difficulty of accurately segmenting tissue data, two annotators tasked with labelling the same image may produce different results. The degree to which human experts agree with each other defines the upper limit of human performance for this task. To understand how the segmentation predictions from Mesmer compared with human performance, we compared Mesmer’s segmentation predictions on four distinct tissue types with predictions from five independent expert human annotators on the same tissues. We evaluated all pairs of human annotators against one another, using one annotator as the “ground truth” and the other as the prediction. We then evaluated Mesmer’s predictions against predictions from each of these five annotators. We detected no significant differences between human-to-human and human-to-Mesmer F1 scores (p=0.93) (**Figure 3d**), suggesting that Mesmer performed on par with human annotators.

To further evaluate Mesmer’s performance relative to humans, we enlisted four pathologists to perform a blinded evaluation of segmentations from the human annotators and Mesmer. Each pathologist was shown a series of paired images, one containing a prediction from Mesmer and the other containing an annotation from a human. The pathologists then rated the relative accuracy while blinded to the origin of each image (**Figure 3e**). When evaluated in aggregate, the pathologists rated Mesmer’s predictions and the expert annotator’s predictions equivalently (**Figure 3f**). Breaking down the evaluation by tissue type, we observed only modest differences in pathologist evaluation. We found one tissue type where Mesmer was rated higher (immune), two tissue types where the evaluation was equivalent (esophagus and breast), and one where the annotators were rated higher (colon) (**Figure 3f**). The single tissue type where Mesmer was not rated as highly suffered from quality issues (see following paragraph), likely underlying the lower score. Taken together, the preceding analyses demonstrate that Mesmer performs whole-cell segmentation with human-level performance. To our knowledge, this is the first time that a cell segmentation algorithm has achieved parity with human segmentations for tissue data.

To finish our performance analysis, we sought to understand Mesmer’s limitations by identifying images where Mesmer produced low-quality cell segmentations. Inaccurately segmented images were characterized by low signal-to-noise ratio, heterogeneous staining, and focus issues (**Figure S3i**). To characterize the impact of each of these factors on model performance, we evaluated model accuracy after blurring, resizing, or adding image noise. While Mesmer was robust to moderate image distortion, performance suffered as the distortions increased in magnitude (**Figure S3j-l**)—as expected, since these manipulations remove information from the images.

### Whole-cell segmentation enables accurate downstream analysis of tissue imaging data

Cell segmentation is the first step for quantitative analysis of tissue imaging data and serves as the foundation for subsequent single-cell analysis. Mesmer’s ability to generate both whole-cell and nuclear segmentation predictions enables new analyses that were difficult to perform with previous segmentation algorithms. One example is predicting the subcellular localization of proteins within cells, which can yield additional information beyond simply quantifying overall intensity. For example, nuclear translocation of transcription factors like NF-κB and IRF3 is informative for understanding signaling dynamics in immune cells^63,64^, while the degree of membrane staining of HER2 is an important feature in the accurate grading and assessment of breast cancer^65^. To explore the accuracy of subcellular signal prediction, we stained breast cancer samples with a panel of phenotyping markers and imaged them with MIBI-TOF^51^ (Methods; **Figure 4a**). We created an integrated multiplexed image analysis pipeline, ark-analysis^66^, that integrates Mesmer’s segmentation predictions with downstream analysis. We used this pipeline to generate nuclear and whole-cell segmentation predictions for each cell in each image. We extracted the compartment-specific expression of each marker in the panel, then employed the same pipeline to extract compartment-specific expression using the ground-truth labels for the same images (Methods). Subcellular localization predictions from Mesmer agreed with those from the ground-truth data (**Figure 4b**). We observed predominantly nuclear expression for known nuclear markers (e.g. Ki67 and HH3) and non-nuclear expression for membrane markers (e.g. E-cadherin and HER2; **Figure 4b**). Thus, Mesmer’s capability for simultaneous nuclear and whole-cell segmentation enables compartment-specific expression to be extracted in an automated fashion.

**Figure 4:**
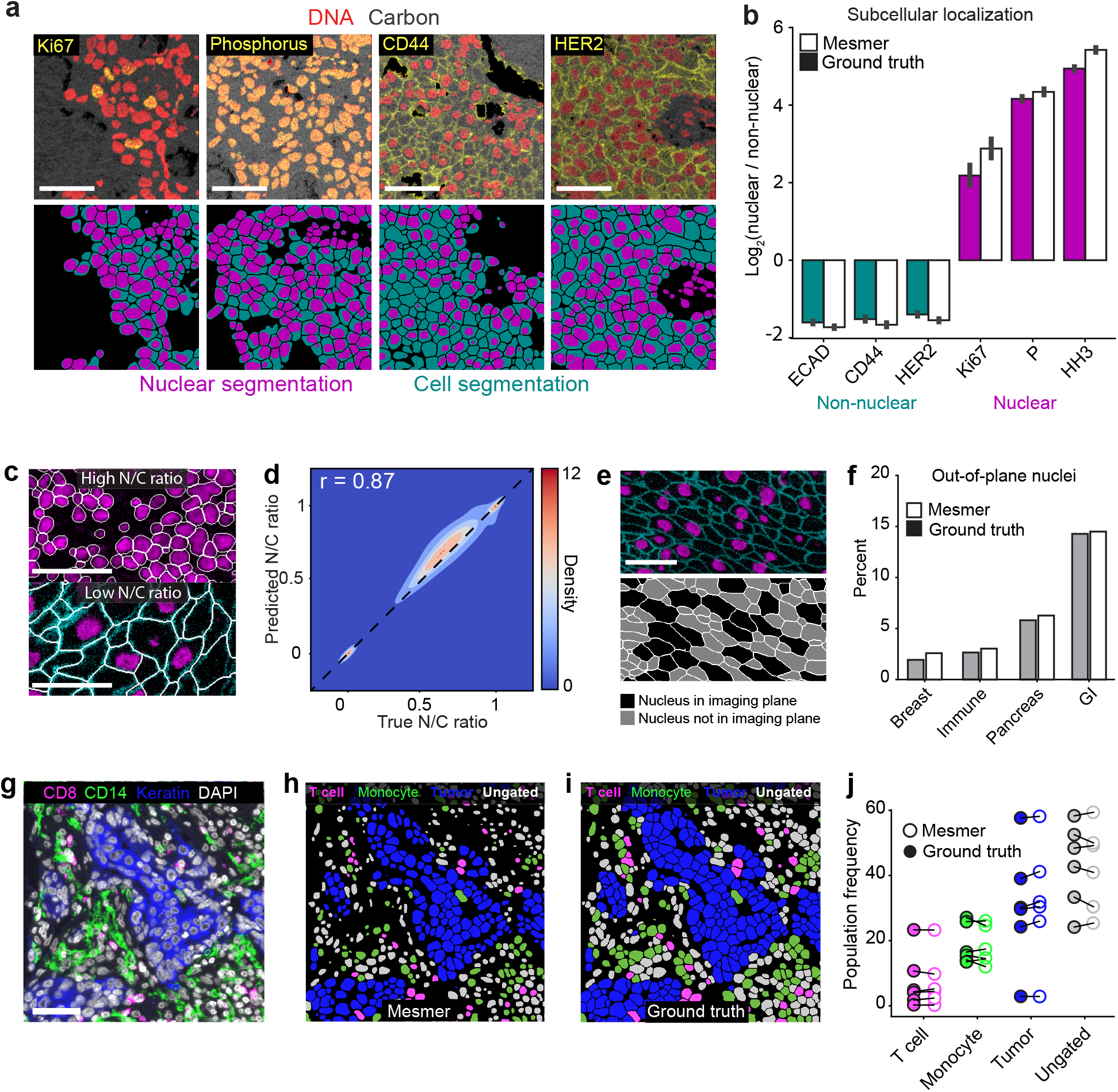
Mesmer enables accurate analysis of multiplex imaging data. **a**, Color overlays showing staining patterns for nuclear and non-nuclear proteins (top), with associated nuclear and whole-cell segmentation predictions (bottom). **b**, Quantification of subcellular localization of the proteins in **a** for predicted and ground-truth segmentations. The agreement between localization for prediction and ground-truth segmentations indicates that Mesmer accurately quantifies protein localization patterns at the single-cell level. **c**, Example image of a tissue with a high N/C ratio (top) and a low N/C ratio (bottom). The N/C ratio is one of several metrics used for quantifying cell morphology (Methods). **d**, A Pearson’s correlation contour plot of the accuracy of N/C ratio predictions across the entire test split of TissueNet demonstrates that Mesmer accurately quantifies cell morphology. **e**, Representative image of a tissue with many nuclei outside the imaging plane (top), along with corresponding segmentations colored by whether the nucleus is or is not in the imaging plane. **f**, Quantification of the number of cells with an out-of-plane nucleus in the predicted and ground-truth segmentations. These cells are detected by Mesmer but would be missed by nuclear segmentation-based methods. GI, gastrointestinal. **g**, Representative image of the expression of multiple informative proteins in a breast cancer sample. **h**, Predicted segmentation colored by cell lineage. **i**, Ground-truth segmentation colored by cell lineage. **j**, Quantification of the frequency of each cell type in the ground-truth and predicted segmentations after gating demonstrates that Mesmer produces accurate cell-type counts. All scale bars are 50 μM.

In addition to subcellular localization, Mesmer also enables the automated analysis of the relationship between each individual nucleus and cell. The nuclear/cytoplasmic ratio is widely used by pathologists to evaluate malignancies; changes in this ratio can have important implications for the speed and aggressiveness with which a tumor grows^67^. To demonstrate that this scoring can be automated, we used Mesmer to generate nuclear and whole-cell segmentations for every cell in the test set of TissueNet. We then computed the nuclear/whole-cell (N/C) ratio, which is conceptually similar to the nuclear/cytoplasm ratio but has higher numerical stability for cells with little cytoplasm (e.g., immune cells; Methods). Mesmer accurately captured cells with low and high N/C ratios (**Figure 4c**), and there was a strong correlation (Pearson’s r = 0.87) between the predicted and ground-truth N/C ratios across all cells in TissueNet (**Figure 4d**).

This analysis identified a subpopulation of cells with an N/C ratio of 0 (**Figure 4e**), indicating that no nucleus was observed in that cell. These cells arise when the imaging plane used to acquire the data captures the cytoplasm, but not the nucleus. We quantified the proportion of cells with an out-of-plane nucleus across the tissue types in TissueNet for both the predicted and ground-truth segmentation labels and found good agreement between predicted and true rates of out-of-plane nuclei (**Figure 4f**). The highest proportion of out-of-plane nuclei occurred in gastrointestinal tissue (**Figure 4f**), presumably due to the elongated nature of the columnar epithelium. Cells with out-of-plane-nuclei are missed by nucleus-based segmentation approaches but are captured by Mesmer.

Cell classification is a common task following segmentation. Inaccuracies in segmentation can lead to substantial bias in the identification and enumeration of the cells present in an image. To benchmark how Mesmer’s predictions impact this process, we analyzed a cohort of breast cancer samples generated with the Vectra platform using our integrated analysis pipeline. Each image was stained with a panel of lineage-defining markers, which we used to classify each cell as either a T cell, monocyte, tumor cell, or ungated (**Figure 4g**). We selected two distinct regions from three patients and generated both predicted and ground-truth segmentations for all of the cells in the image. We classified all cells from the predicted (**Figure 4h**) and ground-truth (**Figure 4i**) segmentations into these categories using the same gating scheme (Methods). Looking at the proportion of cells in each image assigned to each category, we observed strong agreement between the two annotations (**Figure 4j**), showing that Mesmer’s segmentation predictions enable accurate classification of the diversity of cells present in these images.

### Lineage-aware segmentation quantifies cell morphology changes during human pregnancy

Our driving goal in developing the initial release of TissueNet was to cover the tissue types and imaging platforms most frequently analyzed and utilized by the imaging community. This advance would enable the development of general-purpose cell-segmentation models for the majority of use cases while requiring the minimum number of imaging channels. While Mesmer achieved this goal, a small number of organs not represented in TissueNet have cells with morphologies that differ substantially from those currently represented in TissueNet. In some of these cases, the complexity of cellular features exceeds what can be accurately captured with only two segmentation channels. Consequently, these out-of-distribution examples will likely pose a challenge to two-channel models trained exclusively on TissueNet.

Here we focused on one such outlier tissue type, the decidua. The decidua is the mucosal membrane of the uterus and undergoes extensive tissue remodeling during pregnancy. During this time, human endometrial stromal cells transition from an elongated morphology with spindle-like projections to a more globular shape in a process known as decidualization^68^. These large cells are accompanied by ameboid-like extravillous trophoblasts, small mature lymphocytes, pleomorphic uterine macrophages, secretory columnar epithelium, and spiral arteries surrounded by concentric, fragmented layers of vascular smooth muscle^69^. The complexity resulting from such a wide variation in cell shape and size is compounded by the tight juxtaposition of these cells with one another and the non-convex geometries that they can assume^70^. Small round cells often nestle within concavities of much larger cells. Additionally, the large size, abundant cytoplasm, and elongated projections of human endometrial stromal cells often manifest in standard four-micron serial tissue sections as oddly shaped cytoplasmic cross sections with out-of-plane nuclei.

This complex morphology makes segmentation challenging when using a single membrane channel, even for an expert annotator (**Figure 5a**, top). However, information about the location and shape of each cell can be attained by including additional markers that are cell-type specific (**Figure 5a**, bottom). These additional markers provide crucial information about cell morphology during model training that is lost when they are combined into a single channel, leading to poor accuracy (data not shown).

**Figure 5:**
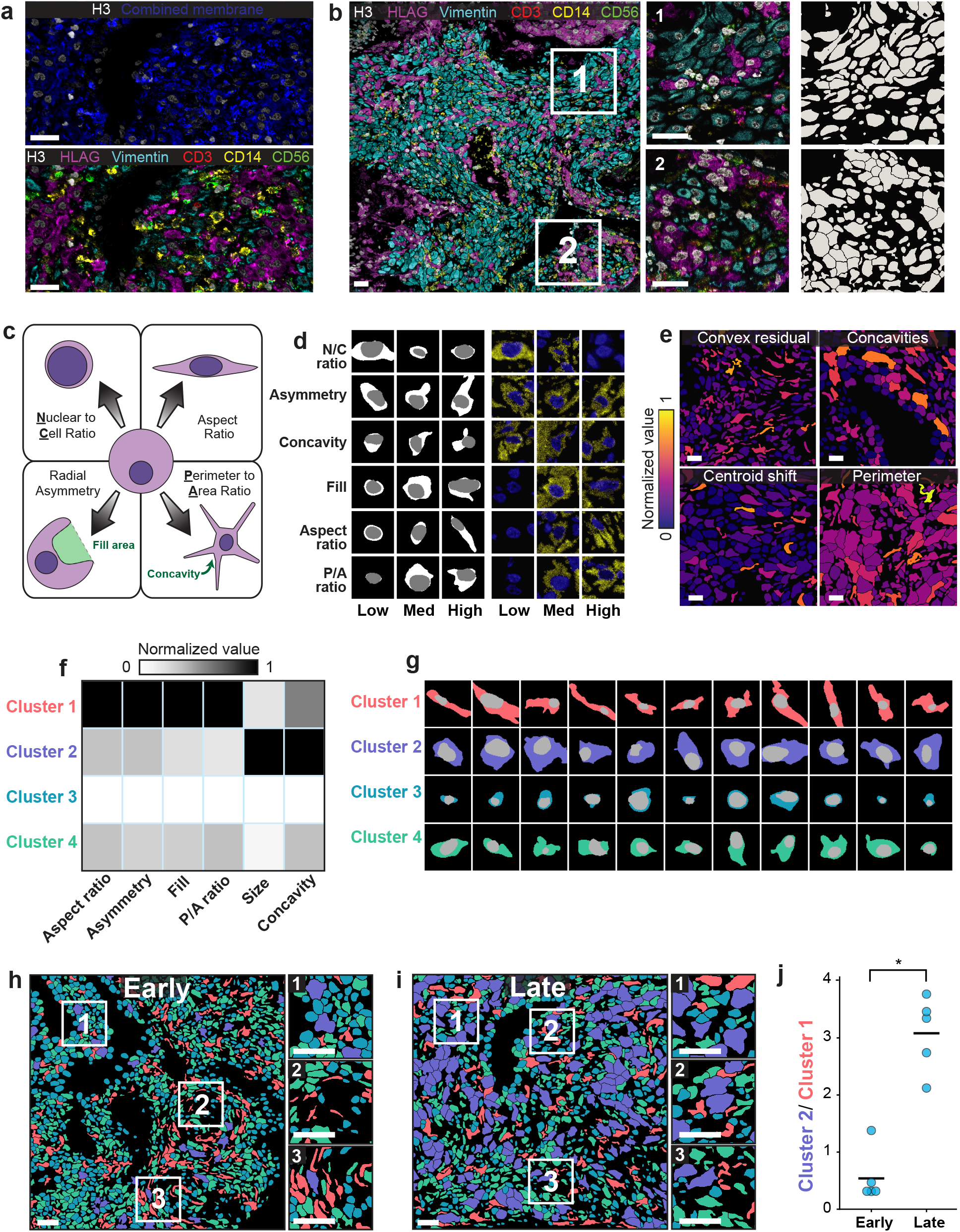
Lineage-aware segmentation yields morphological profiling of cells in the decidua during human pregnancy. **a**, Color overlay showcasing the challenge of distinguishing cells with only a single combined membrane channel (top), paired with a version of the same image containing all six channels used for lineage-aware segmentation (bottom). **b**, Representative image of the diverse morphology of cell types in the human decidua (left), along with insets (right) with corresponding segmentation predictions. **c**, Diagram illustrating the morphology metrics that we defined to enable automated extraction of cell-shape parameters (Methods). **d**, Predicted segmentations (left) placing cells on a spectrum from low to high for each morphology metric, along with the corresponding imaging data for those cells (right). **e**, Cell segmentations in four representative images colored by morphology metrics demonstrate the accurate quantification of diverse features. **f**, Heatmap of inputs to *k*-means clustering used to identify distinct cell populations based on cell morphology. **g**, Example cells belonging to each cluster illustrate the morphological differences between cells belonging to each cluster. **h**,**i** Representative images of maternal decidua in early (**h**) and late (**i**) gestation, with segmentations colored by cluster. **j**, Quantification of the ratio between cluster 2 and cluster 1 cells in early pregnancy versus late-pregnancy samples. Cluster 2 cells become more prominent in the later time point while cluster 1 cells become rarer. *p<0.05, two-sample t-test. All scale bars are 50 μM.

We predicted that by generating training data from images that include multiple lineage markers, it would be possible to train an algorithm to use those additional channels to guide segmentation decisions. We therefore used MIBI-TOF to generate a multiplexed imaging dataset from the human decidua (Greenbaum et al., manuscript in preparation). The antibody panel used to create this dataset had six lineage markers: HH3, Vimentin, HLAG, CD3, CD14, and CD56. We used DeepCell Label^43^ to generate training data from a subset of the images in this cohort. We used DeepCell Label’s capability to simultaneously display six distinct channels at once to generate lineage-aware ground-truth segmentations. We modified our PanopticNet architecture to accept six channels of input data, and trained a model using this dataset (Methods). We then used this lineage-specific segmentation model to replace the TissueNet-based model in Mesmer. The resulting lineage-aware segmentation pipeline accurately performed whole-cell segmentation, despite the complex cell morphologies in these images (**Figure 5b**).

We used this lineage-aware segmentation pipeline to quantify morphological changes of cells in the decidua over time. We first defined a series of morphological metrics to capture the diversity of cell shapes in this dataset (**Figure 5c**; Methods). *N/C ratio* compares the size of the nucleus to the size of the whole cell. *Asymmetry* measures how much a cell deviates from a radially symmetric shape. *Concavities* are indentations in each cell that result in a non-convex shape, and the *fill* metric quantifies the total area that these features occupy. The *aspect ratio* (major axis/minor axis) quantifies cell elongation, while the *P/A ratio* compares the perimeter and area of each cell (perimeter^2^/area). Manual inspection of cells with low, medium, or high values for each of these metrics demonstrated accurate assignment of cells within each category (**Figure 5d**). We then created an automated pipeline that computes each of these metrics for every cell in an image^66^. We applied our pipeline to this dataset and found that these metrics captured key morphological features of the cell shapes that we observed (**Figure 5e**).

To integrate the morphological features that we extracted, we performed *k*-means clustering on the morphology profiles that we collected for every cell (Methods). We identified four distinct clusters (**Figure 5f, g**). Cluster 1 cells were elongated with projections while Cluster 2 cells were typically larger and biased toward non-convex shapes. Cluster 3 cells were small and circular, while Cluster 4 cells exhibited intermediate levels of all features. In order to determine how these cellular morphologies changed over time in the human decidua, we divided the samples into two groups based on age: early (6-8 weeks) and late (16-18 weeks) gestational age. Coloring each cell by its cluster identity highlighted the difference in cell morphology between the two gestational age groups (**Figure 5h, i**). We observed an abundance of cluster 1 cells (e.g., elongated) in the early time point and an abundance of cluster 2 cells (e.g. large and globular) at the late timepoint (**Figure 5j**). This shift likely reflects the morphological transformation undergone by maternal stromal cells during decidualization^71^. During this process, maternal stromal cells transition from an elongated morphology with spindle-like projections to a more globular secretory morphology in the first and second trimesters of pregnancy. Our analysis demonstrates that whole-cell segmentation can make cell morphology a quantitative observable, bridging the historical knowledge of pathologists and modern multiplexed imaging methods.

### DeepCell supports community-wide deployment of Mesmer

Adoption of deep learning methods in the life sciences has been relatively slow, in part due to difficulties in deploying deep learning models. Accessing these methods requires significant computational expertise, and the need for specialized hardware poses a considerable barrier for many labs. Several recent tools have sought to overcome this barrier with a variety of software-engineering approaches, including browser-based software (ImJoy^41^), Google Colab (ZeroCostDL4Mic^42^), a centralized web portal (NucleAIzer^29^, Cellpose^28^, DeepCell^39^), and ImageJ plugins (StarDist^60^, DeepCell^39^).

To spur the adoption of deep learning methods for cellular image analysis in the life sciences, we have created DeepCell, a collection of linked, open-source software libraries that contain tools for generating training data (DeepCell Label^43^), training deep learning models (deepcell-tf^72^), and deploying those models at scale in the cloud (DeepCell Kiosk^39^). While there are existing tools designed to solve each of these problems in isolation, DeepCell seeks to provide an integrated, full-stack solution to the data, model, and deployment needs for using deep learning models for biological discovery. The source code for these software libraries is available for non-profit use through a modified Apache license.

We used DeepCell to make Mesmer accessible to the broader biological imaging community, with two distinct deployment solutions based on the volume of data that must be processed (**Figure 6**). The first solution is geared toward moderate amounts of data (<10^3^ 1-megapixel images) and centers around our web portal https://deepcell.org, which hosts the full Mesmer pipeline. Users can access Mesmer through this web portal directly or submit images through plugins that we have made for ImageJ^21^ and QuPath^73^, two popular image-analysis tools. This web portal is served by a scalable backend (created by DeepCell Kiosk^39^), which dynamically adjusts the server’s computational resources to match the volume of data being submitted. This strategy increases computational resources to support large volumes of data during times of high demand, while reducing these resources during times of low demand to reduce costs.

**Figure 6:**
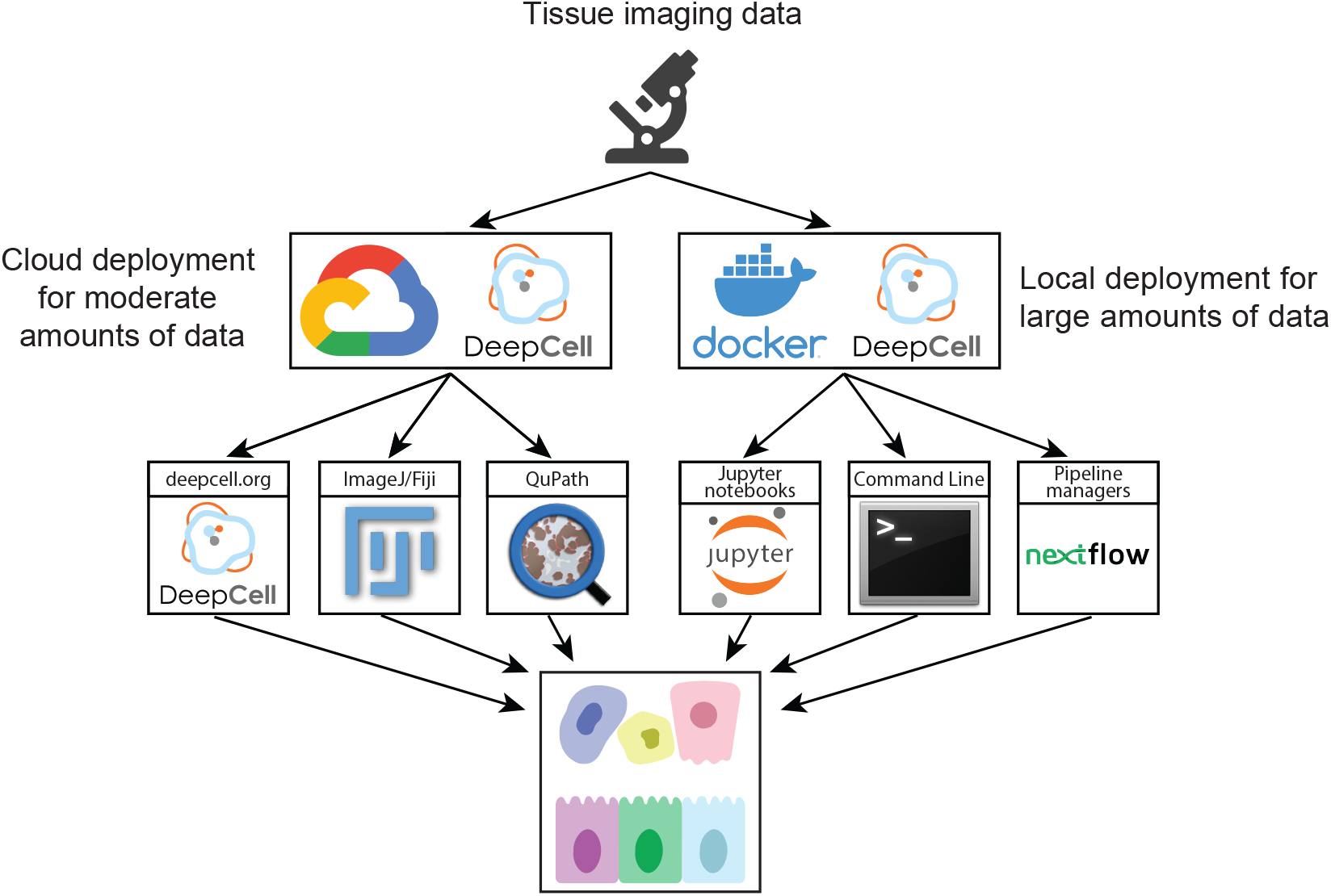
Cloud-native and on-premise software facilitates deployment of Mesmer. A centralized web server, https://deepcell.org, hosts a version of the Mesmer pipeline. Users with moderate amounts of data (<10^3^ 1-megapixel images) to process can access this pipeline through a web portal. Alternatively, users can use ImageJ and QuPath plugins that submit data to the https://deepcell.org web server and receive the results. We have also created a containerized version of Mesmer that is compatible with existing workflow managers, so that users with larger amounts of data (>10^3^ 1-megapixel images) to process can benefit from our work.

The second deployment solution is targeted toward users with larger volumes of data (>10^3^ 1-megapixel images) and who need more control over the execution of the described algorithm. For these users, we provide a Docker image of deepcell-tf that contains the full Mesmer pipeline. The image generates a Docker container locally on a user’s computational infrastructure and can be installed with a one-line command. This container can be used to launch an interactive Jupyter Notebook that processes data with Mesmer. The container can also be configured as an executable, making it possible to integrate Mesmer within existing image-analysis workflows. In addition, we have developed a software package specifically for analyzing multiplexed imaging data, ark-analysis^66^, that integrates cloud-based segmentation predictions with downstream analysis and visualization. We believe that this combination of cloud and local deployment options will make Mesmer accessible to the entire imaging community.

## Discussion

Cell segmentation has been a major bottleneck for tissue imaging community, as prior methods^19,22–24^ required extensive manual curation and parameter tuning to produce usable results. These shortcomings are substantial, and our own experience has shown that they can lead to months-long delays in analysis. Mesmer provides a single unified solution to cell segmentation for the most widely used fluorescence and mass spectrometry imaging platforms in a user-friendly format. Mesmer achieves human-level accuracy across a variety of tissues and imaging modalities while requiring no manual parameter tuning from the end user. To make Mesmer widely available, we created cloud-based and local software solutions that position users of all backgrounds to generate accurate predictions for their data. We anticipate that this pipeline will accelerate discovery in the life sciences by lowering a significant technical barrier to wider adoption of multiplexed imaging platforms. Further, Mesmer’s speed and scalability will facilitate the analysis of the large volumes of multiplexed imaging data currently being generated by consortia around the world.

Along with Mesmer, we present the initial release of TissueNet, the first comprehensive cell segmentation dataset for tissue images. TissueNet contains paired nuclear and whole-cell annotations for >1 million cells from nine organs and six imaging platforms. Prior tissue datasets^16,28^ were not large enough to train accurate whole-cell segmentation models. As a result, previous efforts to generate accurate tissue-segmentation models have focused on nuclear segmentation^27,29,38^, and nearly all previous benchmarking has therefore been limited to the evaluation of nuclear-segmentation models^27,29,60^. TissueNet will enable these valuable efforts to move from nuclear segmentation to whole-cell segmentation, facilitating the development and benchmarking of a new class of tissue-segmentation algorithms. To expand upon this initial release of TissueNet, we are currently constructing a seamless mechanism for investigators to add their own annotated data. Future releases of TissueNet will likely include higher-dimensional data from a wider diversity of tissue types, imaging techniques, and species.

Constructing TissueNet required a new process for generating manual annotations. Traditionally, segmentation datasets have been created by dedicated manual curation with expert annotators. This approach can produce accurate segmentations, but it does not scale to large dataset sizes, as the time required for each new annotation remains constant over the course of the entire project. Our human-in-the-loop approach allowed annotators to benefit from improvements in model accuracy. Because annotators only needed to correct the mistakes made by the model, not annotate every cell in each image, annotation time was linked to model performance. As model performance improved, annotation time decreased, facilitating the creation of substantially larger datasets than was feasible or economical before. We demonstrated the potential of this approach in constructing TissueNet, but we believe this to be the first example of many useful applications of human-in-the-loop dataset construction. We anticipate that there will be an increasing need for large, annotated datasets of biological images, and we believe that constructing these datasets will only be tractable by taking a human-in-the-loop approach.

While Mesmer produced accurate segmentations for a diversity of tissue types, out-of-distribution tissues like decidua with complex cell morphologies remain challenging. For multiplexed images of these tissues, lineage-aware segmentation offers a path forward. Along these lines, we demonstrated the viability of this approach by developing a custom six-channel model for accurate segmentation of the human decidua. While this lineage-aware model was limited to our specific dataset, we believe that it serves as a promising proof of principle for general-purpose, lineage-aware segmentation models. Further, the data annotation, training, and deployment solutions that we have created as part of DeepCell will help to accelerate these subsequent efforts.

While this work is a significant step forward, some challenges remain, the first being the lack of a standardized, cross-compatible antibody panel for cell segmentation. Development of such a panel would be a major advance and would synergize with the work presented here. Whole-cell segmentation in 3D is another challenge that will become more prominent as imaging throughput increases to allow routine collection of such datasets. Existing deep learning approaches for 3D instance segmentation are promising^74^, but a 3D equivalent of TissueNet to power future models currently does not exist. Our work can serve as a starting point for these efforts, as it yields accurate prediction in 2D slices of tissues (**Figure S4**). Now that accurate cell segmentation is available to the entire community, it will be exciting to see the novel insights delivered by the diversity of data currently being generated.

## Methods

### Creating TissueNet

#### Human-in-the-loop annotation in the crowd with DeepCell Label

Labeling multiplexed imaging data presents a unique software engineering challenge. Labeling software needs to allow users to view multiple channels at once, so that they can use all available information to identify cell and nuclear boundaries. This software also needs specialized labeling operations to enable efficient labeling of densely packed fields of cells. Further operations are needed for creating labels from scratch and for editing existing labels; the latter is a key requirement for our human-in-the-loop framework. The final constraint is that this software needs to be browser based, which is essential for crowdsourcing. While existing software packages address specific aspects of these challenges, to date none have met all of the necessary requirements for human-in-the-loop data annotation of multiplexed image data.

To meet this challenge, we previously developed DeepCell Label^43^, a software package by which humans and algorithms collaboratively create and correct annotations for biological images. DeepCell Label consists of a frontend, which enables users to visualize and interact with images and labels (**Figure S1a**), and a backend, which serves images and labels (stored in cloud buckets) to the frontend (**Figure S1b**). This backend is built on Elastic Beanstalk, a scalable web framework that allows our application to scale as the number of users increases. This scalability enables multiple users to work on the same collection of data at the same time while maintaining responsiveness. A database keeps track of user access and stores key metadata involved in the annotation process. The DeepCell Label software is available athttps://github.com/vanvalenlab/deepcell-label

Because DeepCell Label is cloud-native, it is compatible with any crowdsourcing platform that supports HTML iframes. We have successfully used two crowdsourcing platforms to perform crowdsourced labeling of multiplexed imaging data with DeepCell Label: Appen (https://appen.com) and Anolytics (https://anolytics.ai). DeepCell Label enables our human-in-the-loop framework to blend expert and novice human annotators to increase the scale of creating dense, pixel-level labels for biological images (**Figure 1**). An example of the instructions provided for the crowd annotators can be found here:https://github.com/vanvalenlab/publication-figures/blob/mesmer_update/2021-Greenwald_Miller_et_al-Mesmer/Example_annotation_instructions.docx

#### Cropping and stitching of labeled images

To support accurate crowd annotation of dense images, we found that supplying smaller image crops led to significantly better crowd performance (data not shown). Large images can be overwhelming for annotators to examine and are difficult to navigate at the high zoom level necessary for accurate pixel-level annotation. These two issues significantly increase the time required to complete each job. To alleviate these issues, we created a pipeline to crop and stitch images as part of the annotation process (**Figure S1c**). Input images are cropped to a predetermined size, generally 270×270 pixels, with an overlap between adjacent crops. We keep track of the necessary metadata for each crop to facilitate stitching the image back together. Each crop is independently annotated, with crops from the same image being randomly assigned to annotators. Following annotation, these crops are stitched back together. Cells at the boundary between crops are merged based on maximum overlap. Once each image has been stitched back together, it is quality-controlled by an internal expert to correct stitching artifacts and remaining errors from the annotator output. The finalized annotations are stored with the corresponding image data in .npz files to facilitate easy loading and manipulation in python.

#### Combining labeled data together for model training

To construct the dataset used for model training, individual .npz files containing annotated images from a single experiment were combined. During this process, the data were randomly split 80%/10%/10% into training, validation, and testing fractions. We applied automated quality control to each image, such as removing cells with area <15 pixels and removing images with <20 cells. Finally, we cropped each image to 256×256 pixels, the input size that the model expects.

#### TissueNet construction

To construct TissueNet, we collected published and unpublished data from numerous tissue imaging labs^44–51^. Each dataset was manually inspected to identify images suitable for model training. To be included, images from each dataset needed to have robust nuclear staining of all cells, as well as membranous/cytoplasmic staining of a substantial subset of the cells. For datasets with multiple potential nuclear and membrane markers, the best marker (high signal-to-noise ratio, ubiquitous expression) was chosen for each cell type. For multiplexed datasets containing more than one high-quality nuclear or membrane marker, these channels were added together (after rescaling) in cases if doing so produced increased coverage of the relevant cell compartment across the cells in the image. Selected images were fed through the human-in-the-loop data pipeline to create the final labeled dataset.

### PanopticNet architecture design

#### PanopticNet architecture

The deep learning models used for segmentation are panoptic feature pyramids. Briefly, these networks consist of a ResNet50^54^ backbone that is connected to a feature pyramid. Prior to entering the backbone model, images are concatenated with a coordinate map. We use backbone layers C3-C5 and pyramid layers P3-P7; for the pyramid layers we use depthwise convolutions for computational efficiency. We attach two semantic segmentation heads to the feature pyramid that perform upsampling to produce predictions the same size as the input image.

#### PanopticNet transforms

We employed a deep learning approach to segmentation that is inspired by previous work^30,59^. For each image we use a deep learning model to predict two transforms of that image’s label image. The first is a prediction whether each pixel belongs to the cell interior, cell boundary, or background. We call this transform a “pixel-wise transform”. The second is a transform that captures the distance of each pixel inside a cell to that cell’s centroid. If the distance of a cell’s pixel to that cell’s centroid is r, then we compute 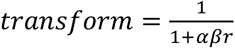 where 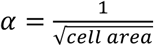 and β is a hyper-parameter that we take to be 1. We call this the “inner distance transform”. One key difference between our formula and the work where this strategy was first proposed^30^ is the introduction of α, which makes this transform relatively indifferent to differences in cell size. We use the softmax loss for the semantic head assigned to the pixel-wise transform and the mean squared error for the semantic head assigned to the inner distance transform. Similar to prior work, we scale the softmax loss by 0.01 to stabilize training.

#### PanopticNet preprocessing

To handle the variation in staining intensity and signal-to-noise ratio across tissue types and imaging platforms, we normalize each image prior to running it through our PanopticNet architecture. We first perform 99.9% scaling to reduce the influence of extremely bright, isolated pixels. We then utilize Contrast Limited Adaptive Histogram Equalization (CLAHE)^56^ to normalize each image to have the same dynamic range.

#### PanopticNet post-processing

The output of the model is two sets of predictions, one for the interior/border/background transform and a second for the inner distance transform. We use marker-based watershed^58^ as a post-processing step to convert these continuous predictions into discrete label images where the pixels belonging to each cell are assigned a unique integer id. To perform this post-processing step, a peak-finding algorithm^75^ is first applied to the prediction image for the inner distance transform to locate the centroid of each cell in the image. These predictions are thresholded at a value of 0.1. The interior class of the prediction image for the pixel-wise transform is thresholded at a value of 0.3. The cell centroid locations and interior pixel prediction image are used as inputs to the marker-based watershed algorithm to produce the final label image. We smooth the transforms with a gaussian filter to eliminate minor variations and perform additional processing that removes holes and all objects with an area <15 pixels from the final prediction.

#### PanopticNet training

All models were trained using the Adam optimizer^76^ with a learning rate of 10^−4^, clipnorm of 0.001, and batch size of 8 images. During training, each image is augmented by performing random flips, rotations, crops, and scaling to increase the diversity of the training dataset. We use 80% of the data for training, 10% for validation, and 10% for testing. We evaluate the loss on the validation dataset after each epoch, and only save the model weights if the loss decreases from the previous value. The test set is only used to evaluate the final trained model. The PanopticNet code is available at: https://github.com/vanvalenlab/deepcell-tf

### PanopticNet architecture evaluation

#### Classifying error types

We previously described a methodology for classifying segmentation error types^37^, which we used here. Briefly, we use linear assignment to find the best match between the cells in the ground truth-labeled data and the predicted cells in the model prediction. We use the intersection over union (IOU) of the cell masks as the criterion for assessing whether two cells match, with thresholds of 0.4 and 0.1 for the cost matrix and IOU overlaps, respectively. If there is a direct, 1-to-1 match between a true cell and a predicted cell, then that cell is classified as accurately segmented. If a true cell has no matches, then it is classified as a false negative, as there is no corresponding predicted cell. If a predicted cell has no matches, then it is classified as false positive, as there is no corresponding true cell. If two predicted cells are assigned to the same true cell, then these three cells are part of a split error. If two true cells are assigned to the same predicted cell, then these three cells are part of a merge error. Finally, if none of the above criteria are satisfied, and there is a complex relationship between multiple true and predicted cells, then we categorize this cell as “other.”

#### Benchmarking model performance

To evaluate model accuracy, we created three random splits of TissueNet, each with a different training, validation, and testing set. These three separate versions of TissueNet were used to train models in triplicate for all model comparisons in this and subsequent sections. Each model was trained using the training and validation splits and evaluated using the corresponding test split, which was never seen during training. We used our error classification framework to calculate the types of errors present in each image in the test split and reported the average and standard deviation across the replicates. All PanopticNet models were trained for 100 epochs with a fixed number of steps per epoch to control for differences in dataset size.

#### Comparison with alternate models and architectures

To evaluate the benefits of the PanopticNet architecture, we compared our approach with several alternative algorithms. These algorithms included Cellpose^28^ (whole-cell), StarDist^60^ (nuclear), RetinaMask^59^ (trained on TissueNet), FeatureNet^16^ (nuclear), and FeatureNet^26^ (trained on TissueNet). All evaluations were performed using the same three splits of TissueNet.

### Mesmer performance analysis

#### Nuclear expansion comparison

To compare Mesmer with the current nuclear-based segmentation approaches listed above, we generated whole-cell labels using Mesmer, as well expanded nuclear labels, for all of the images in the test set. Nuclear expansion predictions were generated from nuclear predictions by applying a morphological dilation with a disk of radius 3 pixels as the structuring element. To characterize the error modes of each approach, we selected predictions that mapped directly to a single ground-truth cell using our error-type classification approach (see *Classifying error types*). Following identification of the corresponding ground-truth cell, we computed the ratio of the predicted cell size to true cell size for each prediction.

#### Specialist model evaluation

To evaluate how a specialist model trained on only a subset of the data compared to a generalist model trained on the entire dataset, we identified the four most common tissue types and four most common imaging platforms in TissueNet. Each of these four tissue types had images from multiple imaging platforms, and each of the four imaging platforms had images from multiple tissue types. For each of the eight specialist models, we identified the images in the training and validation split that belonged to that class and used that subset for model training. We then evaluated the trained specialist model on the data in the test split that belonged to that class and compared this performance to the generalist model evaluated on the same portion of the test split.

#### Dataset size evaluation

To evaluate how training dataset size impacts model accuracy, we divided TissueNet into bins of increasing size. Each bin of increasing size contained all of the data from the previous bins, with new data added. This strategy ensures that each bin is a superset of the previous bin, rather than each bin being a random draw from the whole dataset. Bins of increasing size were generated for the training and validation splits, while holding the test split constant. We trained models on the progressively larger bins and evaluated all models on the same complete test set.

#### Inter-annotator agreement

To determine the degree to which annotators agreed with one another, we recruited five expert annotators (lab members or PhD students) to annotate the same set of images. For each of the four images, all five annotators generated segmentations from scratch, without using model predictions. We also generated model predictions for these same four images, which were not included in the training data. We then computed the F1 score between all pairs of annotators, as well as between each annotator and the model.

#### Pathologist evaluation

To evaluate the relative accuracy of Mesmer and human annotators, we enlisted four board-certified pathologists to evaluate segmentation accuracy. Each pathologist was shown pairs of images, where one image contained the segmentation predictions from Mesmer and the other contained the segmentation prediction from one of our expert annotators. We selected 13 random crops from each of the four images. Each crop was displayed to each pathologist twice, with the same Mesmer prediction each time, but matched to a different expert annotator prediction, for a total of 104 comparisons.

#### Image distortion quantification

To determine how image quality impacts model performance, we systematically degraded images in the test set and assessed the corresponding decrease in F1 score. To simulate out-of-focus images, we performed a gaussian blur with increasing sigma. The blurred images were then passed through the model to generate predictions. To determine how image resolution impacts model performance, we downsampled each image to represent low-resolution data. We then upsampled back to the original size and ran the upsampled images through the model. To simulate low signal-to-noise ratio and high background staining, we added uniform random noise of increasing magnitude to each pixel. The noise-corrupted images were passed through the model to generate predictions.

### Analyzing multiplexed imaging datasets

#### Generating subcellular segmentation predictions

We created a custom analysis pipeline that integrates nuclear and whole-cell segmentation predictions. This pipeline takes as inputs the predictions from Mesmer of each cell and each nucleus in the image. We first link each cell mask with its corresponding nuclear mask using maximum overlap, splitting nuclei that are larger than their corresponding cell. We use these linked masks to extract the counts per compartment for all channels of imaging data. The counts for each marker within each compartment are summed and normalized by cell area. Our multiplex image analysis pipeline is available at: https://github.com/angelolab/ark-analysis

We used this pipeline to compute the subcellular localization of a panel of phenotypic markers with known profiles. We stained a tissue microarray of ductal carcinoma in-situ samples, imaged them with MIBI-TOF, and ran the above pipeline. For each channel, we selected fields of view in which the marker showed clear expression, and computed the localization within each cell, after removing the bottom 20% lowly expressing cells within each marker. We performed the same procedure using the ground-truth labels generated by the human annotators and used the computed localization from the true labels to assess the accuracy of our predictions.

#### Computing nucleus to whole-cell ratio (N/C)

Traditionally, the nuclear to cytoplasm ratio assessed by pathologists is the ratio between the area of the nucleus and the area of the cytoplasm^67^. However, as a quantitative measure, this formulation runs into issues with division by zero for immune or stromal cells that have no detectable cytoplasm. To alleviate this issue, we instead use the nucleus to whole-cell ratio (N/C), which uses the whole-cell area rather than the cytoplasm’s area. The nuclear and whole-cell areas are always greater than zero, thus avoiding division by zero; this leads to more stable estimates while maintaining the same qualitative interpretation. Cells with high N/C ratios have larger nuclei relative to their overall cell size, and cells with low N/C ratios have smaller nuclei relative to their overall cell size.

#### Evaluating accuracy of N/C ratio predictions

To determine the accuracy of our N/C ratio predictions, we ran Mesmer on the entire test split of TissueNet. We computed the nuclear and cell predictions for each cell in the image. For each cell, we computed the N/C ratio by first matching each nucleus to its corresponding cell, and then calculated the ratios of their respective areas. We reported the Pearson correlation between the true N/C and predicted N/C for all predicted cells with a direct match in the ground-truth data.

#### Assessing frequency of out-of-plane nuclei

Multiplexed imaging platforms analyze tissue slices that represent a 2D cut through a 3D structure. As a result, sometimes the nucleus of a given cell is not captured in the image plane/tissue section, whereas the rest of the cell is. Given that Mesmer is trained to separately identify nuclei and cells, cells whose nucleus is out of the imaging plane can still be identified and segmented. To determine the frequency of this occurrence, and to validate that these predictions were not merely segmentation artifacts, we compared the incidence of cells with an out-of-plane nucleus in the ground truth data and Mesmer predictions from the TissueNet test set. For each ground truth or predicted cell, we identified the corresponding nucleus. Cells without a matching nucleus were classified as out of plane.

#### Quantifying accuracy of cell lineage frequency

To determine how the accuracy of Mesmer’s segmentation predictions influenced downstream quantification of cell type, we analyzed a cohort of breast cancer samples acquired on the Vectra platform. We selected two fields of view each from three unique patients. Each patient’s sample was stained with DAPI, CD8, CD14, and Pan-Keratin to identify the major cell subpopulations. We generated segmentation predictions for all images with Mesmer, as well as ground-truth labels with our human-in-the-loop pipeline. We then extracted the counts of each marker in each cell and used hierarchical gating to define cell populations. Thresholds for gating were determined by manual inspection of the histogram for size-normalized counts of each marker. We used the same thresholds for the ground-truth and predicted segmentations. For each field of view, we quantified the proportion of cells assigned to each lineage, and then linked the predictions for a given cell type in a given field of view to the corresponding ground-truth proportion.

### Decidual cell morphology

#### Training a six-channel Mesmer model

To establish the potential of a model that takes in multiple lineage markers, we retrained the PanopticNet architecture within Mesmer. We generated images of human decidua using the MIBI-TOF platform (Greenbaum et al., manuscript in prep), and stained the samples with a panel of markers to define the cell lineages present. We manually generated whole-cell segmentation labels for 15 of these images, using HH3 to define the nucleus, and CD3, CD14, CD56, HLAG, and vimentin to define the shape of the cells in the image. We modified the model architecture to accept six channels of input data, and trained it using the settings described above.

#### Generating cell morphology information

To quantify the range of cell shapes and morphologies present in the image data, we created an automated pipeline that extracts key features from each cell segmentation in an image. We extract morphological information using the regionprops function in the scikit-image^75^ library. We use the following default features from regionprops: area, perimeter, centroid, convex area, equivalent diameter, convex image, and major axis length. These features are transformed as described below to create the selected morphology metrics. Our analysis pipeline is available at: https://github.com/angelolab/ark-analysis

Many of the metrics relate to the difference between the cell shape and the corresponding convex hull. A convex hull for a given segmentation is defined as the smallest possible convex shape that completely contains the cell shape. For shapes that are already convex and do not have any concave angles, the convex hull and the cell are equivalent. For shapes that do have concavities, the convex hull fills in these areas.

In addition to the N/C ratio, we computed the following five morphology metrics:

- *Asymmetry:*the distance between the centroid of the convex hull and the centroid of the cell, normalized by the square root of the area of the cell. The centroid of the convex hull is far from the centroid of the cell when extra mass is added to the convex hull in an imbalanced fashion, indicating that the original cell was not symmetrical.
- *Concavities:*the number of concavities present in each cell. We only include concavities that have an area of at least 10 pixels^2^ and a perimeter-to-area ratio < 60 to avoid counting very small deviations from convexity. This approach summarizes how many unique indentations and divots are present in each cell.
- *Fill:*the difference in area between the convex hull and the cell, normalized by the area of the convex hull. This ratio is effectively the proportion of the convex hull that was newly added and quantifies the fraction of the cell that is composed of divots and indentations.
- *Aspect ratio:*the ratio between the major axis length and the equivalent diameter. Major axis length is the length of the major axis of an ellipse with the same moments as the original cell and serves as a proxy for the length of the longest diagonal of the cell. Equivalent diameter is the diameter of a circle with the same area as the cell. The ratio of these two quantities gives an estimate of cell elongation.
- *Perimeter to area ratio:*The ratio between the perimeter squared of the cell and the area of the cell. We use perimeter squared rather than perimeter itself for better consistency across cell sizes.

#### Identifying morphological clusters

We classified the cells based only on the above five morphology metrics, which we computed for the images in the decidua cohort. We first normalized each metric independently, and then performed *k*-means clustering with *k*=4. We plotted the mean value of metric in each cluster to identify the features that separated them from one another and performed hierarchical clustering on the resulting output.

### Model deployment

#### DeepCell Kiosk: A scalable, cloud-based solution for hosting deep learning models

We previously described the construction of DeepCell Kiosk, our cloud-based deployment system^39^. This software dynamically adjusts the amount of compute resources needed at any one time to match demand; since the load on the server is quite low most of the time, this strategy delivers economical hosting of the web portal for community use. When demand increases, compute resources are automatically increased. The Kiosk is available at: https://github.com/vanvalenlab/kiosk-console.

#### Generating predictions from Mesmer using cloud deployments

To facilitate quick and easy access to Mesmer, we used the Kiosk to generate a number of easy ways to predict cell segmentation. We created a web portal that allows anyone to upload their data and instantly receive results. This web-based interface facilitates point-and-click upload and download of results, with no installation required. We have also created plugins for ImageJ^21^ and QuPath^73^ that automatically send data to the Kiosk and return predictions to the user. These predictions can then be used within ImageJ or QuPath for downstream analyses of interest. Detailed tutorials and documentation can be found at https://github.com/vanvalenlab/intro-to-deepcell.

#### Generating predictions from Mesmer using local deployments

Although cloud deployment offers a fast, intuitive way for users with little computational experience to generate predictions, it offers less fine-grained control over the input and output parameters. Further, web portals are not ideal for integration with existing image processing workflows. For users with more computational expertise, we have created local deployments of Mesmer to facilitate future model development and integration with existing workflows. To facilitate training and model development, we provide example Jupyter and Colab notebooks. For integration with existing computational workflows, we provide a command line interface and docker container. Finally, we have also made our own open-source multiplex image analysis pipeline available for users who want an end-to-end solution for segmenting, quantifying, and analyzing image data. A guide showing users how to use these resources is available at https://github.com/vanvalenlab/intro-to-deepcell.

### Software

This project would not have been possible without numerous open-source python packages. These packages include jupyter^77^ keras^78^, matplotlib^79^, numpy^80^, pandas^81^, scikit-image^75^, scikit-learn^82^, seaborn^83^, tensorflow^84^, and xarray^85^. Specific versions for each package can be found at https://github.com/vanvalenlab/deepcell-tf/blob/master/requirements.txt.

## Acknowledgments

We thank Long Cai, Katy Borner, Matt Thomson, Steve Quake, and Markus Covert for interesting discussions; Sean Bendall, David Glass, and Erin McCaffrey for feedback on the manuscript; Roshan Angoshtari, Graham Barlow, Bernd Bodenmiller, Christopher Carey, Robert Coffey, Alea Delmastro, Colt Egelston, Michal Hoppe, Hartland Jackson, Anand Jeyasekharan, Sizun Jiang, Youn Kim, Erin McCaffrey, Eliot McKinley, Michael Nelson, Siok-Bian Ng, Gary Nolan, Sanjay Patel, Yanfen Peng, Darci Philips, Rumana Rashid, Scott Rodig, Sandro Santagata, Christian Schuerch, Daniel Schulz, Diana Simons, Peter Sorger, Jason Weirather, and Yuan Yuan for providing imaging data for TissueNet; the crowd annotators who powered our human-in-the-loop pipeline; and all patients who donated samples for this study.

## Authorship Contributions

N.F.G., L.K., M.A., and D.V.V. conceived the project. E.M. and D.V.V. conceived the human-in-the-loop approach. L.K. and M.A. conceived the whole-cell segmentation approach. G.M., T.D., E.M., W.G., and D.V.V. developed DeepCell Label. G.M., N.F.G., E.M., I.C., W.G., and D.V.V. developed the human-in-the-loop pipeline. M.S., C.P., W.G., and D.V.V. developed PanopticNets. W.G., N.F.G., and D.V.V. developed model training software. C.P. and W.G. developed cloud deployment. M.S., S.C., W.G., and D.V.V. developed metrics software. W.G. developed plug-ins. N.F.G., A.Kong, A.Kagel, J.S., and O.B-T. developed the multiplex image analysis pipeline. A.Kagel and G.M. developed the pathologist evaluation software. N.F.G., G.M., and T.H. supervised training data creation. N.F.G., C.C.F., B.M., K.L., M.F., G.C., Z.A., J.M. and S.W. performed quality control on the training data. E.S., S.G., and T.R. generated MIBI-TOF data for morphological analyses. N.F.G., W.G., and D.V.V. trained the models. N.F.G., W.G., G.M., and D.V.V. performed data analysis. N.F.G., G.M., M.A., and D.V.V. wrote the manuscript. M.A. and D.V.V. supervised the project. All authors provided feedback on the manuscript.

## Data and software availability

All data will be made available at datasets.deepcell.org upon publication in a peer-reviewed journal.

All software for dataset construction, model training, deployment, and analysis is available on our github page https://github.com/vanvalenlab/intro-to-deepcell. All code to generate the figures in this paper is available at https://github.com/vanvalenlab/publication-figures/tree/master/2021-Greenwald_Miller_et_al-Mesmer

