## Supplementary Figures for "Whole-cell segmentation of tissue images with human-level performance using large-scale data annotation and deep learning"

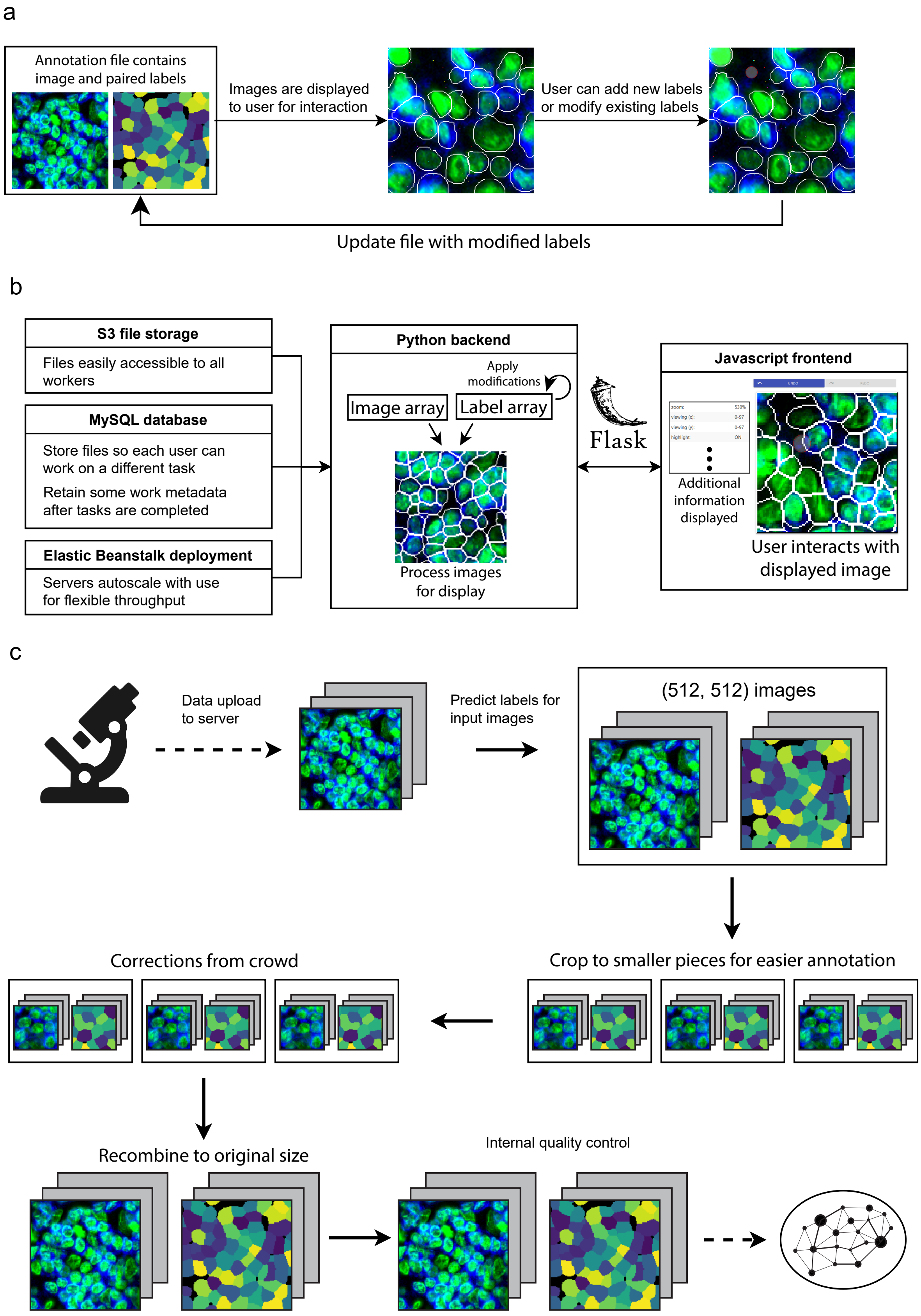

a
Annotation file contains image and paired labels
Images are displayed to user for interaction
User can add new labels or modify existing labels
Update file with modified labels
b
S3 file storage
Files easily accessible to all workers
MySQL database
Store files so each user can work on a different task
Retain some work metadata after tasks are completed
Elastic Beanstalk deployment
Servers autoscale with use for flexible throughput
Python backend
Apply modifications
Image array
Label array
Process images for display
Flask
Javascript frontend
Additional information displayed
User interacts with displayed image
c
Data upload to server
Predict labels for input images
(512, 512) images
Crop to smaller pieces for easier annotation
Corrections from crowd
Recombine to original size
Internal quality control

**Figure S1**. **a**, How multichannel images are represented and edited in DeepCell Label. **b,** Scalable backend for DeepCell Label that dynamically adjusts required resources based on usage, allowing concurrent annotators to work in parallel. **c**, Human-in-the-loop workflow diagram. Images are uploaded to the server, run through Mesmer to make predictions, and cropped to facilitate error correction. These crops are sent to the crowd to be corrected, stitched back together, run through quality control to ensure accuracy, and used to train an updated model.

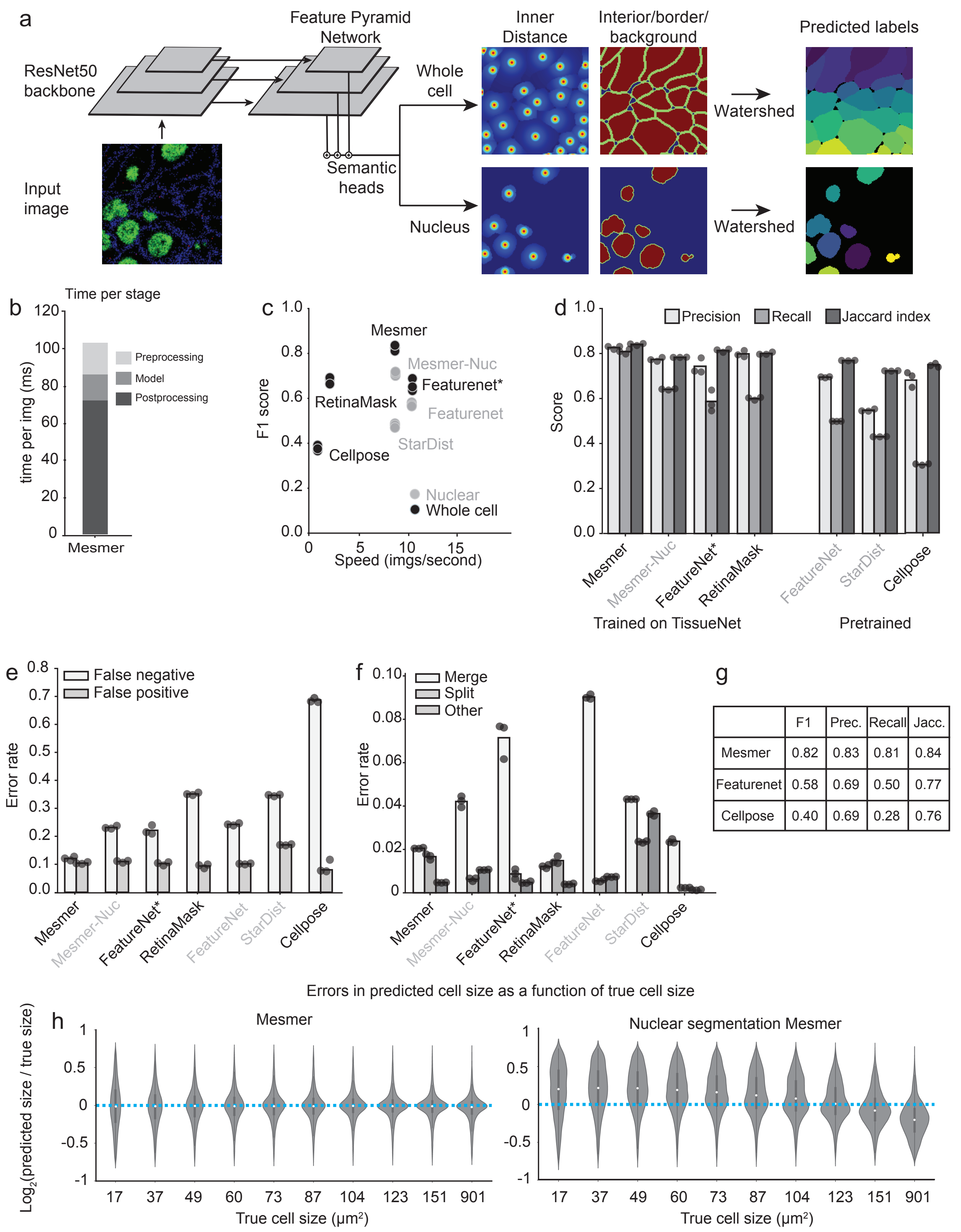


| | F1 | Prec. | Recall | Jacc. |
|---|---|---|---|---|
| Mesmer | 0.82 | 0.83 | 0.81 | 0.84 |
| Featurenet | 0.58 | 0.69 | 0.50 | 0.77 |
| Cellpose | 0.40 | 0.69 | 0.28 | 0.76 |

**Figure S2**. **a,** PanopticNet architecture. Images are fed into a ResNet50 backbone coupled to a feature pyramid network. Two semantic heads produce pixel-level predictions. The first head predicts whether each pixel belongs to the interior, border, or background of a cell, while the second head predicts the center of each cell. **b,** Relative proportion of preprocessing, inference, and post-processing time in PanopticNet architecture. **c**, Speed versus accuracy comparison between Mesmer and previously published models demonstrates the superior performance of Mesmer. **d,** Evaluation of precision, recall, and Jaccard index for Mesmer and previously published models shows that Mesmer outperforms previously published models. **e**,**f** Breakdown of most prevalent error types (**e**) and less prevalent error types (**f**) for Mesmer and previously published models illustrates Mesmer's advantages over previous approaches. **g**, Summary of TissueNet accuracy for Mesmer and selected models to facilitate future benchmarking efforts. **h**, Comparison of the size distribution of prediction errors for Mesmer (left) with nuclear segmentation followed by expansion (right) shows that Mesmer's predictions are unbiased.

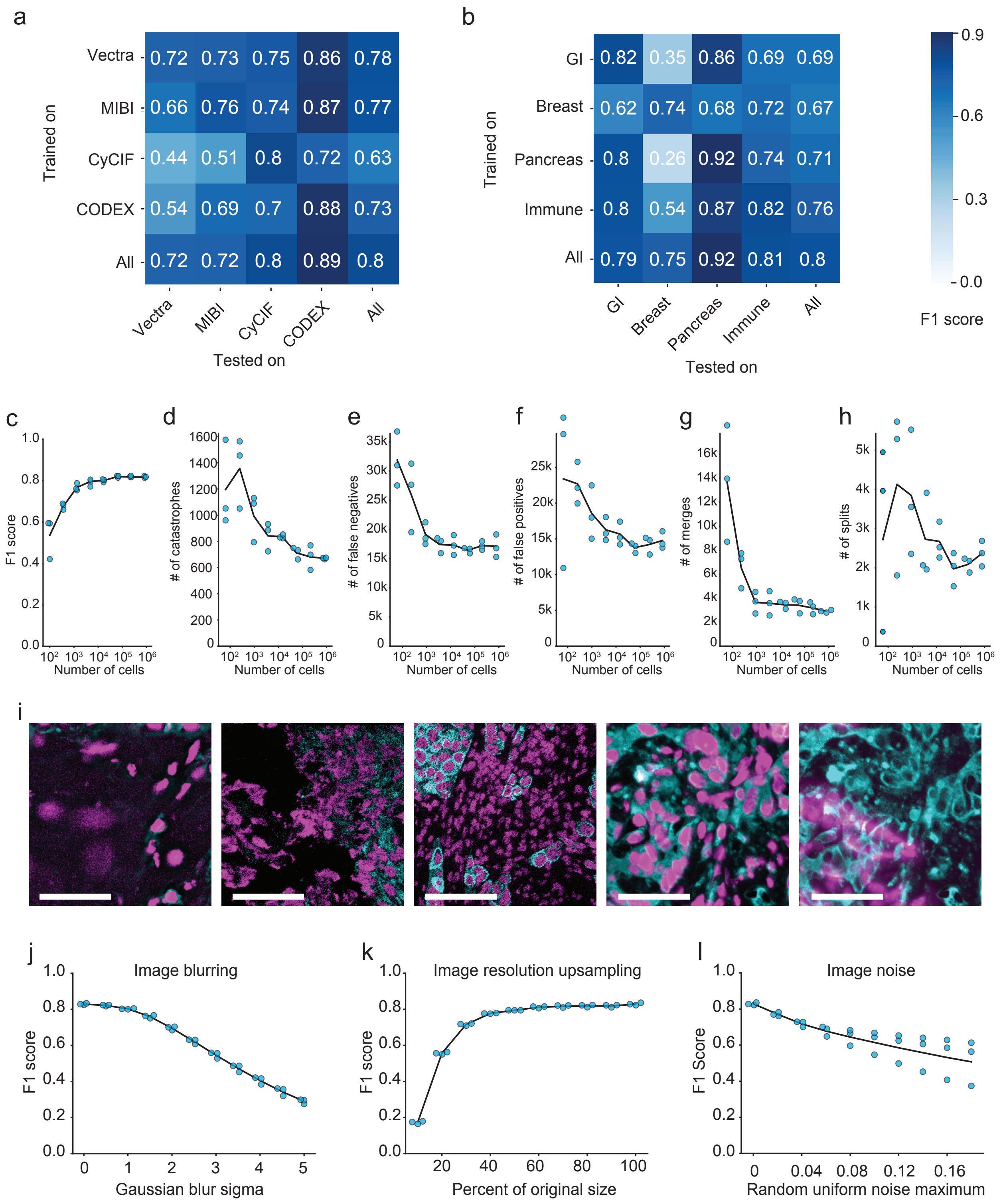


**Figure S3**. **a**, Accuracy of specialist models trained on each platform type (rows) and evaluated on data from other platform types (columns) indicates good agreement within immunofluorescence and mass spectrometry-based methods, but not across distinct methods. **b**, Accuracy of specialist models trained on each tissue type (rows) and evaluated on data from other tissue types (columns) demonstrates that models trained on only a single tissue type do not generalize as well to other tissue types. **c**, Quantification of F1 score as a function of the size of the dataset used for training. **d-h**, Quantification of individual error types as a function of the size of the dataset used for training. **i**, Representative images where Mesmer accuracy was poor, as determined by the image specific F1 score. **j**, Impact of image blurring on model accuracy. **k**, Impact of image downsampling and then upsampling on model accuracy. **l**, Impact of adding random noise to image on model accuracy. All scale bars are 50 μM.

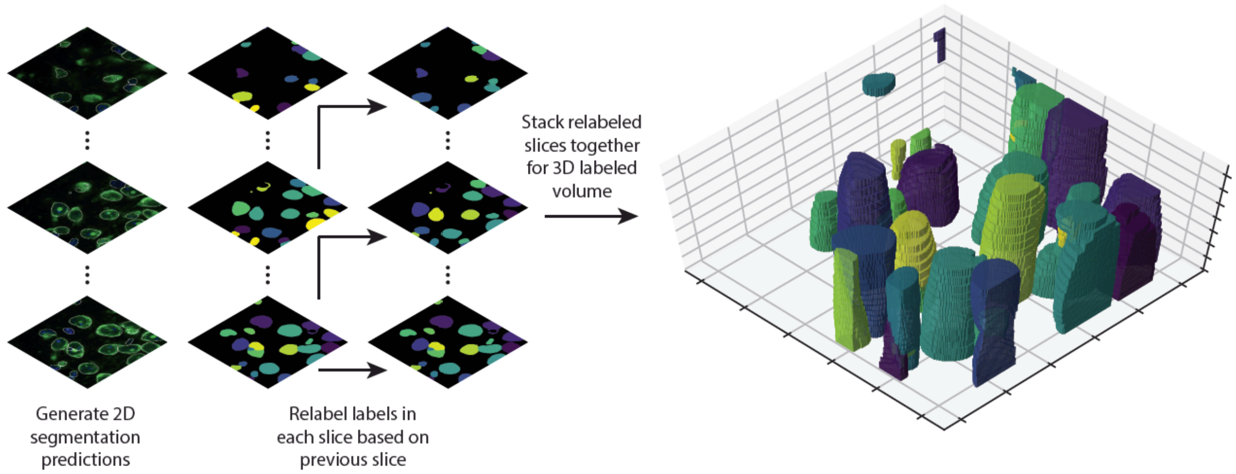


**Figure S4**. Proof of principle for using Mesmer's segmentation predictions to generate 3D segmentations. A z-stack of 3D data is fed to Mesmer, which generates separate 2D predictions for each slice. We computationally link the segmentations predictions from each slice to form 3D objects. This approach can form the basis for human-in-the-loop construction of training data for 3D models.
